# Evolutionary design of optimal surface topographies for biomaterials

**DOI:** 10.1101/2020.05.29.123422

**Authors:** Aliaksei Vasilevich, Aurélie Carlier, David A. Winkler, Shantanu Singh, Jan de Boer

## Abstract

Natural evolution tackles optimization by producing many genetic variants and exposing these variants to selective pressure, resulting in the survival of the fittest. We use high throughput screening of large libraries of materials with differing surface topographies to probe the interactions of implantable device coatings with cells and tissues. However, the vast size of possible parameter design space precludes a brute force approach to screening all topographical possibilities. Here, we took inspiration from Nature to optimize materials surface topographies using evolutionary algorithms. We show that successive cycles of material design, production, fitness assessment, selection, and mutation results in optimization of biomaterials designs. Starting from a small selection of topographically designed surfaces that upregulate expression of an osteogenic marker, we used genetic crossover and random mutagenesis to generate new generations of topographies.

## Introduction

The use of implantable medical devices to treat an increasing variety of human pathologies is growing rapidly ^1^. These devices can support organ function, (e.g. pacemakers), partially replace the organ (e.g. hip implants), or help recover tissue function (e.g. coronary stents). The choice of materials used in the implant is crucial, as its properties determine tissue responses such as inflammation and fibrosis. Poor materials choices can result in implant failure like late stent thrombosis ^2^ or tumour formation induced by breast implants ^3^. The physical and chemical properties of a biomaterial define its multidimensional design space ^4 5^. Materials may have hundreds of properties so design spaces that are generated by combinatorial variation of these properties can be truly enormous (∼10^100^, an essentially infinite number of materials). Very small regions of these vast design spaces can be explored experimentally using high throughput screening of hundreds to thousands of different chemistries ^6^, topographies ^7^, and hydrogels, ^8^ in an array format, screened for a desired biological response. Carefully use of design of experiments (DoA) methods enables these data to represent larger regions of design space than brute force screening alone, and machine learning (ML) methods can interpolate between experimentally measured data points. This has led to the identification of biomaterials that could not be anticipated. For example, we identified a surface topography that supports osteogenesis ^9^. Hook *et al*. discovered specific polymer chemistries with low pathogen attachment properties in three sequential screenings, where materials for subsequent steps were formulated from the best examples from the previous steps ^6^. However, considering the size of the design space, there is a negligible chance that the biomaterial properties we and others found are globally optimal. Quantitative data obtained from screens can be used to explore design space and find correlations between material design and biological activity using ML and other statistical methods ^10,11^. For example, using this approach we were able to predict topography design promoting stem cell proliferation ^12^ and Kholodovych et al. were able to predict clinically relevant responses for diverse polymer designs ^13^. However, the predictive power of ML models is limited to the domain of applicability defined by the properties of materials used to train the model, limiting its scope to discover conceptually different materials that lie well outside the domain. To address this severe limitation in the discovery of improved, novel biomaterial we adopted an evolutionary materials design approach using genetic algorithms (GA) ^14,15^.

Genetic algorithms, inspired by biological evolution, are efficient methods of exploring extremely large parameter spaces. They require material to be represented by a ‘genome’ (a mathematical representation of the physicochemical or processing parameters of a material). ^16^ By analogy with biological evolution, GAs start with a population of biomaterials (existing lead compounds or random materials) with a range of material properties represented their ‘genes’. GAs apply evolutionary pressure in the form of a fitness function (e.g. desired cell response) to generate successive populations of materials by applying genetic crossover, point mutation, or elitism operators to the fittest members of prior populations of materials. Individuals or solutions that best fit a specific environment have a better chance of survival and reproduction. The whole process is repeated until the best solution is found, no fitter solution is found, or a specified number of evolutionary cycles is reached (usually constrained by time or financial resource limitations). As with biological evolution, this evolution pressure generates locally optimal, but nonetheless novel and useful solutions.

Genetic algorithms have previously been used to optimize materials (see the recent review by Le et al. ^17^). For instance, Collins et al. used GAs to optimize functional groups of metal-organic frameworks to increase CO_2_ uptake ^18^. Grand Canonical Monte Carlo calculations were used to calculate the fitness properties (CO_2_ uptake, surface area, and parasitic energy), based on the molecular structure of the compounds. 1.65 trillion structures were analyzed *in silico* and led to the identification and experimental testing of 141 new compounds with increased CO_2_ uptake. In biomaterials research, GAs were employed to optimize the composition of titanium alloys for dental applications ^19^, and to optimize alginate scaffolds for tissue replacement ^20^. In both cases, calculated elastic properties were used as fitness functions.

While ML models of structure-property relationships can be used as surrogate fitness functions, providing the fastest and most convenient fitness assessment for evolutionary optimization ^21^, experimentally measured fitness properties essential to create the data on which the ML models can be trained. For example, Bawazer et al. measured droplet parameters to optimize droplet formation for microfluidics devices ^22^.

In this work, we propose applying cell-based assays as fitness functions for biomaterials optimization using GAs. To assess whether GAs can be used to optimize the bioactive properties of materials we used an existing data set measuring the relationship between the expression of the alkaline phosphatase (ALP, a biomarker of osteogenic differentiation) in mesenchymal stem cell (MSCs) and surface topography. We performed a screen to identify which topographies enhance osteogenesis in titanium implants ^9^. The initial population consisted of 2176 randomly generated surface topographies ^7^. In the future, we will use GAs to optimize surface topographies using multiple cycles of manufacture, biological screening, and mutation to generated increasingly fitter examples of surface topographies. Here we describe the initial step, the evolutionary design of a second generation TopoChip.

## Results and discussion

### Encoding and selecting parents in evolutionary cycles

To generate the initial pool of topographies (described later), and produce genetic mutations of topographies, we converted the information about topography design from a set of design parameters reported in our original TopoChip article ^7^ into a ‘topography genes’. Since no literature exists on the optimal representation of topographies as genes, we developed two different approaches and compared them in subsequent experiments. The first approach is *pixel based* and represents each topographical feature (the base unit of design) as a binary image with a size of 200 × 200 pixels (see Figure 1A). This can be presented as a 2D matrix, populated with 0s and 1s, where 1 corresponds to the pillars of material and 0 means no pillars. The gene is created by flattening this two-dimensional matrix into one dimension by arranging stacks of rows with 0 and 1 into a single row (Figure 1 A). Thus, we converted all topographies on the original TopoChip into genes, one gene representing one surface design.

**Figure 1.**
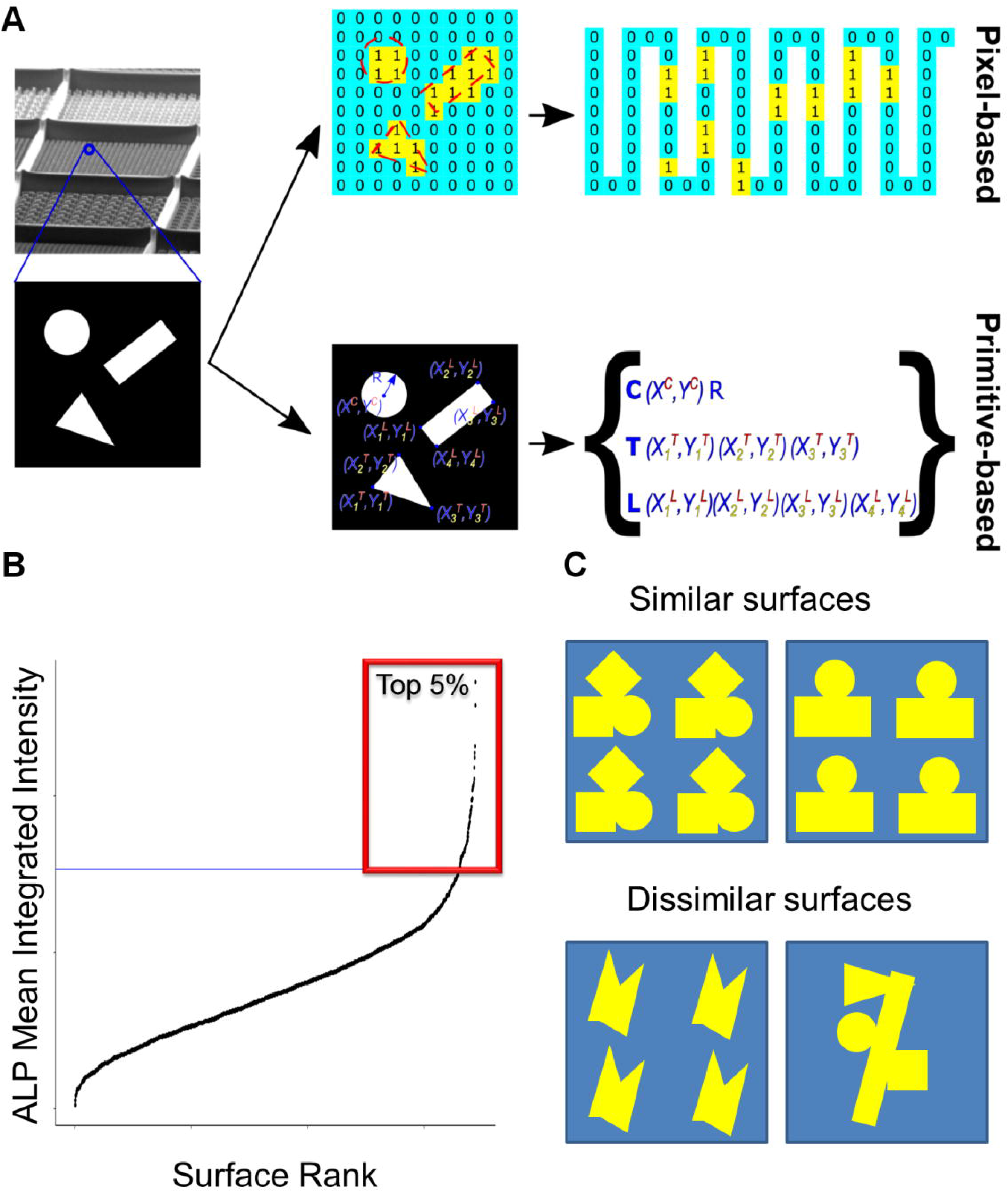
Encoding and selecting parents for cross breeding. **A.** Conversion of surface topography to a gene. Pixel-based approach: the topographical feature can be presented as a 2D matrix, populated with 0s and 1s, where 1 corresponds to the pillars of material and 0 means no pillars. The gene is created by flattening this two-dimensional matrix into one dimension by arranging stacks of rows with 0 and 1 into one row. Primitive-based approach: the gene is defined by primitives of three types (triangle (T), circle(C) or line (L)), the XY positions of lines and triangles corners or XY position of the circle centre and its radius. **B.** Selection of top 5% surface topographies based on ALP Mean Integrated Intensity. **C.** Schematic representation of surfaces that will be similar or dissimilar based on Pearson correlation.

The second approach is *primitive-based* and uses the design parameters which created the first TopoChip. This gene is defined by domains, similar to protein domains encoded by parts of a gene. In this case, each domain encodes a surface geometry, and we chose the three types of primitives (triangle (T), circle (C) or line (L) ^7^). The gene was constructed from the relative XY position of lines or triangle corners, and the center XY position and radius (R) for circles (Figure 1A).

To initiate the evolutionary experiment, we selected 81 parent topographies (top 5% based on their induction of ALP expression) from a pool of 2176 TopoChip topographies ^9^ (Figure 1 B). This list and ranking were based on a previous study in which we exposed bone marrow mesenchymal stem cells to titanium-coated TopoChips and measured the intensity of ALP. These topography ‘parents’ were used to generate millions of diverse progeny topographies using genetic mutation methods. The strategy used to select how the mutation operators are applied to the parents is important as it determines the genetic diversity of progeny. We assessed which of seven selection algorithms (Roulette, Random, Tournament, NSGA2 ^23^, Best, Worst, SPEA2 ^24^) gave the highest genetic diversity in the offspring. Some algorithms (for instance NSGA2) use a similarity score based on Pearson correlation of the genes and ALP expression, others are completely random (Random). Figure 1C explains how the algorithm distinguishes similar and dissimilar surfaces based on Pearson correlation. This defines the computational procedure to align pairs of parents for subsequent mutation.

Breeding and mutation were performed over multiple cycles. In each cycle, groups of 10 parents were selected from an initial pool of 81 parent surfaces. These were combined to generate 10×10 parent pairs, plus the 10 original parents (elitism operator), a total of 110 topographies to be assessed for fitness.

### Application of modification operators to the fittest populations

After parent selection, genetic diversity is created by crossover and point mutations. Parent selection, crossover and mutation are schematically presented in Figure 2 A and 2B. Crossover involves exchanging a substantial block of genes between two selected parents. We implemented eight crossover algorithms, with increasing genetic diversity, for both pixel- and primitive-based parent genomes.

**Figure 2.**
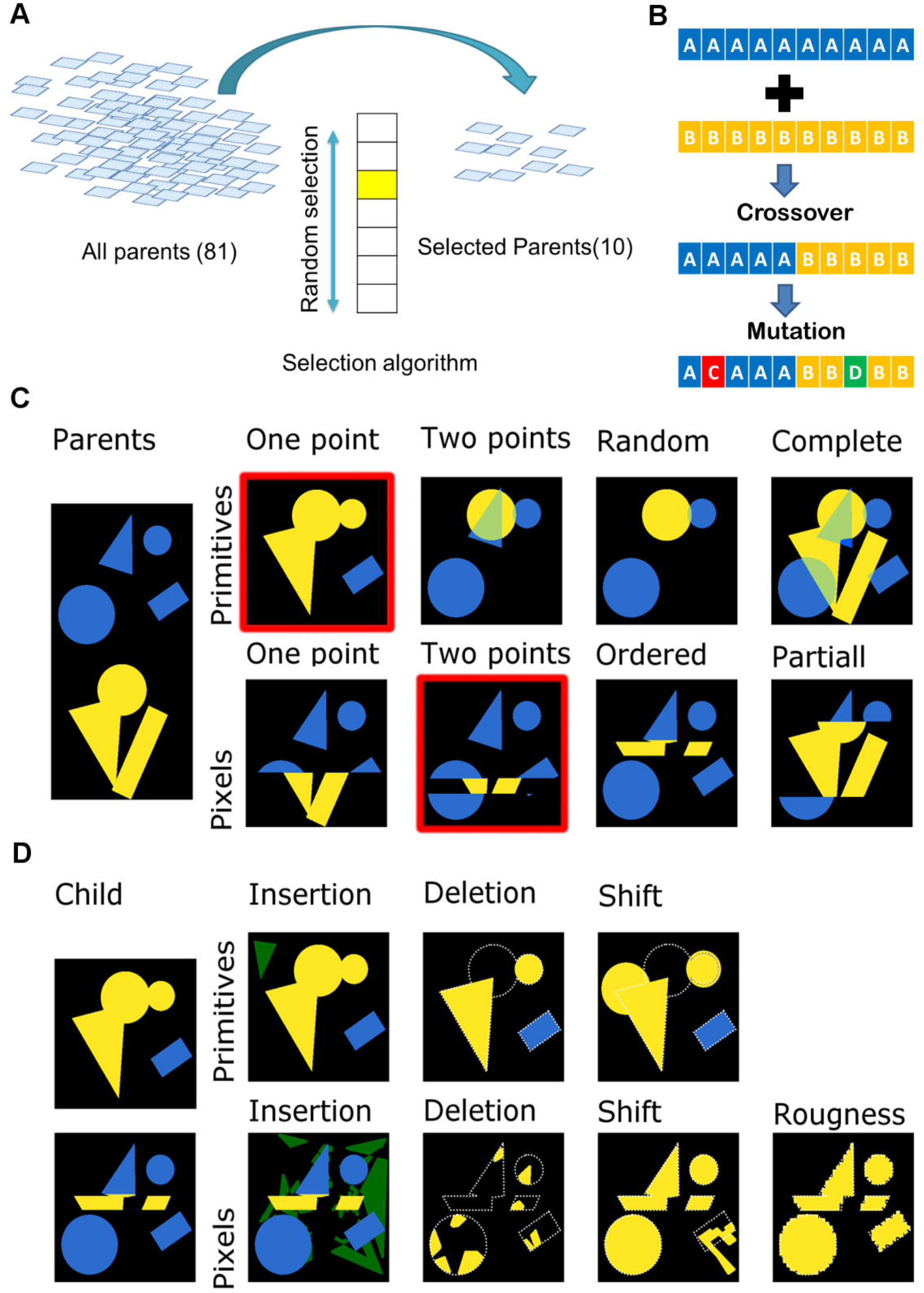
Selecting, recombining and mutating the genetic information of the parents. **A.** Schematic representation of surface selection flow at the beginning of each cycle. **B.** Schematic representation of crossover and mutation algorithms. **C.** Example of different crossover algorithms. Colours (yellow and blue) represent parts from different parents. Surfaces with red outline were selected further to demonstrate mutation algorithms. **D.** Example of different mutation algorithms. Green colour corresponds to newly inserted elements of the design; white outline corresponds to deleted parts.

Figure 2C displays an overview of the effect of different crossover operations on the design of topographies. Visual observation showed that the effect of crossover is clearly different between pixel- and pattern-based representations. Primitive-based representation creates patterns that were similar to the parents, while pixel-based crossover introduced new elements into the topographies compared to the original TopoChip.

After crossover, random mutagenesis operations were applied, introducing a stochastic element to designs. We randomly changed the shapes of the elements, their size (mutation, polygon), partially deleted elements (deletion) or introduced new elements (insertion of circles, triangles, and rectangles). For pixel-based representation, we allowed the insertion of a polygon with a random shape and introduced roughness to the pattern (Figure 2D). Our GA mutation procedure allowed a surface to undergo zero, one, two or three different types of mutations subsequently.

Mutation rate is also a very important parameter. High mutation rates explore larger regions of phenotypic space. This can move topographical designs away from regions where good ALP expression induction occurs, but can also discover distant, novel regions of design space that may also have good APL induction properties. Because we do not know the optimal mutation rate to drive evolution, we tested <20% and <50% mutation rate. Numbers were chosen based on visual observation of the effect of mutation rate on the appearance of the progeny (data not shown). The overall flow of genetic perturbations on topographies is schematically shown in Figure 3A.

**Figure 3.**
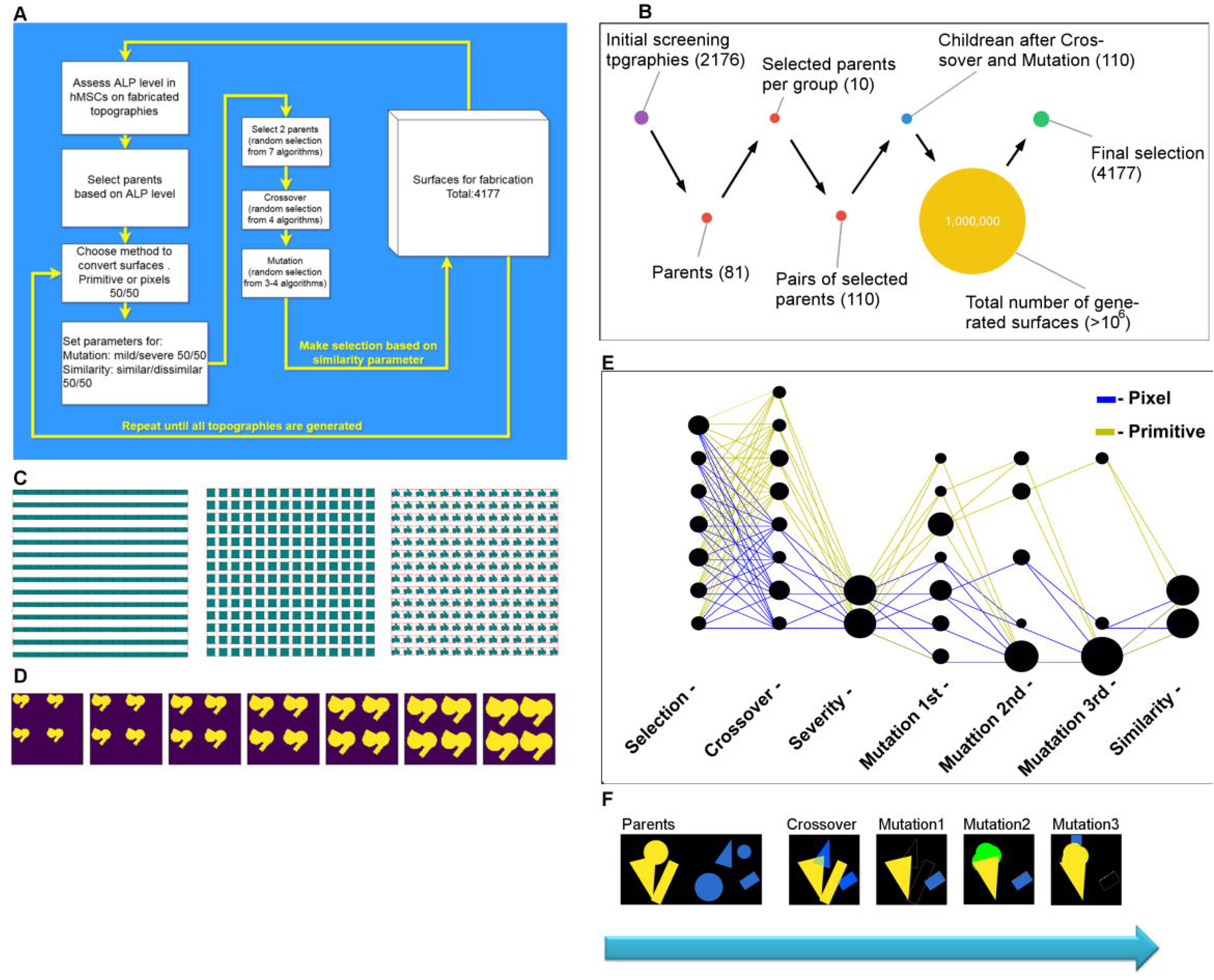
The flow of Genetic Perturbations. **A.** Schematic representation of the genetic perturbation flow. **B.** Schematic representation of surface numbers in every step of genetic perturbation. **C.** Examples of control surfaces. **D.** Different sizes of the ALP hit surface. **E.** The flow of genetic perturbations with the second generation of the GTopoChip1. Every bubble corresponds to a single function or parameter that belongs to step specified on X-axis. The size of the bubble represents the number of surfaces that were generated with that particular function or parameter. **F.** Example of surface generation: after parental crossover one child has 3 consecutive mutation steps. Yellow and blue colours correspond to the parents, green colour represents inserted elements of the design, and white outline represents removed elements of the design.

Each set of 10 parents produced 110 progeny (extra 10 by applying elitism) (see Figure 3B). We used a filter based on similarity measurement in which a pixel-based Pearson correlation coefficient was calculated for each new design against already selected new designs. In the final selection, half of the new surfaces were dissimilar (Pearson correlation index below 0.5) and another half similar (Pearson correlation index below 0.9) to each other. In this way, we produced about one million offspring topographical designs from which 4177 selected for a second generation TopoChip (Figure 3B) to form genetically modified TopoChip (GTopoChip1).

### Types of topographical surfaces

GTopoChip1 has five surfaces types; (1) the 81 original parents, in this way we introduced the elitism operator that allows to keep best surface designs from the previous generations (2) 4177 newly generated surfaces that were perturbed through either pixel-based or (3) primitive-based gene representation as described above, (4) five flat control surfaces, and (5) 31 control surfaces with a simple design in triplicate (Figure 3C). The latter was included to investigate the efficacy of simpler topographies in which surfaces were based on simple gradients or incremental differences in distance between features. This is a commonly-adopted ‘spike-in’ strategy used by many groups working on variants of surface topography. To create simple design surfaces, we took 3 types of patterns, a line, square, and a hit topography from the original TopoChip screen that induced ALP expression and bone bonding in vivo. With the lines and rectangles, we created a gradient of sizes between 4–16 μm in 1 μm steps. 4 μm was the smallest feature that could be produced, and 20 μm was a reasonable maximum feature size. The ALP hit surface was generated in 7 size variants (Figure 3D).

### Exploration of pattern design on GTopoChip1

Figure 3E shows the sequence from parent selection to inclusion in the final design for the 4177 new topographies on the GTopoChip1. The size of the node illustrates the relative number of times the specific node was chosen. The degree of mutation and similarity were pre-selected and are therefore of similar size, Pixel-based and primitive-based algorithms shared the same selection algorithms, while the crossover algorithms were different for each. As Figure 3B shows, most surfaces had only one mutation; a few topographies received 3 consecutive mutations. Figure 3F shows an example of surfaces generated using a crossover and all 3 types of mutations sequentially.

The ALP expression and topographical similarity (based on Pearson correlation) were plotted for the 81 parents as described in Figure 4A. The figure shows the number of children per parent as a bubble chart. Interestingly, topographies that created the most offspring generated diverse ALP induction and had low topographical similarity. This likely reflects the fact that similarity measurements and ALP expression were used in the selection step. Parent selection resulted from 7 randomly selected algorithms. We observed that they are not equally represented on GTopoChip1. In fact, “Random” and “Worst” selection algorithms were the most frequently used among the primitive based and pixel-based gene representations of the surfaces, respectively (Figure 4B). Despite the fact that all algorithms were selected by uniformly distributed random sampling (i.e. equal probability of each algorithm being selected), filtering steps that excluded surfaces with similarity above a specified threshold most likely skewed this distribution, and algorithms that contributed to the generation of unique surfaces were chosen more frequently. Possibly, Worst and Random selection algorithms created more dissimilar surfaces, explaining their more frequent selection.

**Figure 4.**
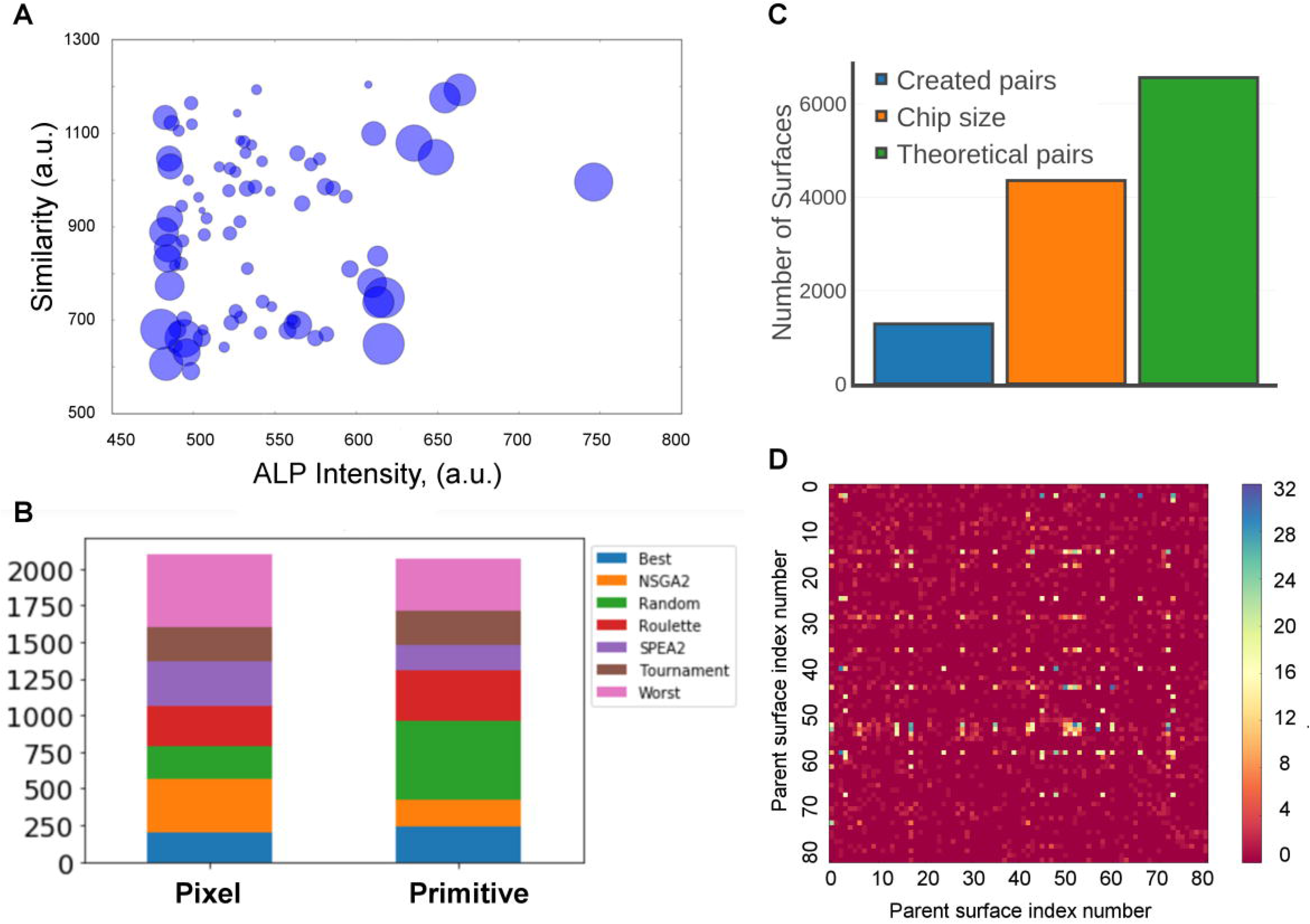
Exploration of Genetically modified TopoChip. **A.** Bubble chart in which the bubble size represents the number of progeny for every parent. Y-axis represents similarity measured with Pearson correlation. The x-axis represents ALP intensity in the initial screen. **B.** The frequency of the selection algorithm for both Pixel-based and primitive based gene representations. C. Quantification of factual, possible and theoretical parents’ pairs. **D.** Quantification of pairs formed by the parent surfaces in GTopoChip1.

We also quantified the total number of unique pairs of parents (Figure 4C). 1284 unique pairs were found from a theoretical maximum of 81×81=6561 that could be formed, (note that the GTopoChip1 can only accommodate 4356 topographies). Some parents contribute more to genetic diversity than others and random mutagenesis contributes significantly to genetic diversity. This is illustrated in Figure 4D, where we plotted all 6561 parents and marked which parent combinations and their number are present on the chip. As can be seen, some parents have bred with a large number of other parents, whereas some barely reproduced. Clearly, some parent combinations were used multiple times and produced multiple offspring because mutations of these combinations increased topographical diversity and fitness. For example, Figure 5 shows all the progeny of the best ALP inducing surfaces from the previous screening. Some progeny are only slightly different from the parent surfaces, while others barely resemble the parents (highlighted by a yellow outline on the figure).

**Figure 5.**
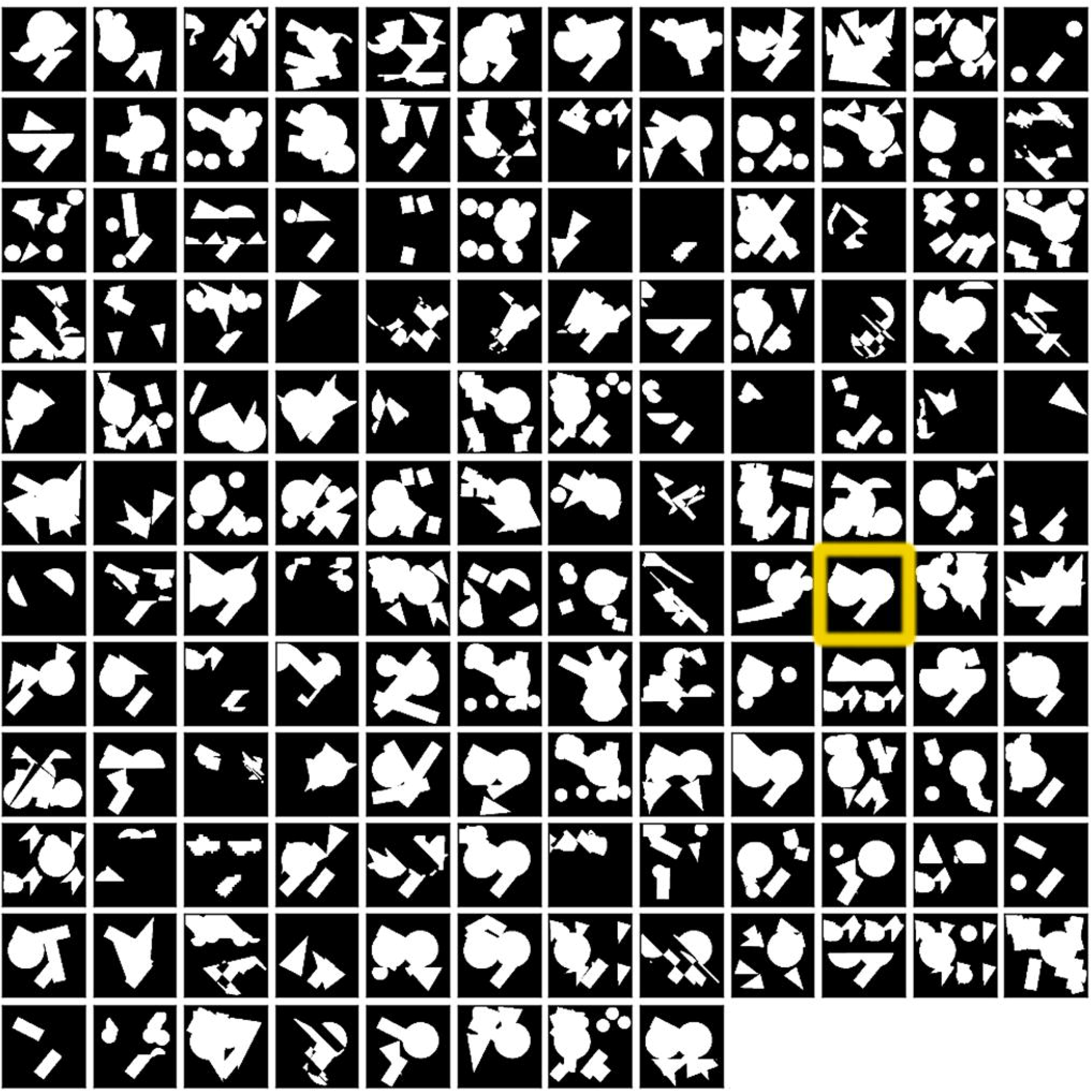
Progeny of the ALP hit surface from the initial screening. Parent surface is enhanced with a yellow outline.

We analyzed similarities and dissimilarities between progeny and parent topographies. All designs were converted to images and then to pixel-based genes. We then used principal components analysis (PCA) to calculate similarity, and visualized GTopoChip1 using the first two principal components (PCs) (Figure 6A). It is clear that the parents do not cover design space evenly; the flat surface occupies one corner in the PCA plot and the control topographies are probing design space only in the first PC. Interestingly, the majority of pixel-based topographies are shifted towards the left, away from the bulk of parents, whereas primitive based surfaces are closer to the parents. This means that when primitive based parents are crossed, their offspring have higher topographical similarity to them than when pixel-based parents are crossed. It is important to realize that it is possible to crossover half a pixel-based circle, but not half a primitive-based circle.

**Figure 6.**
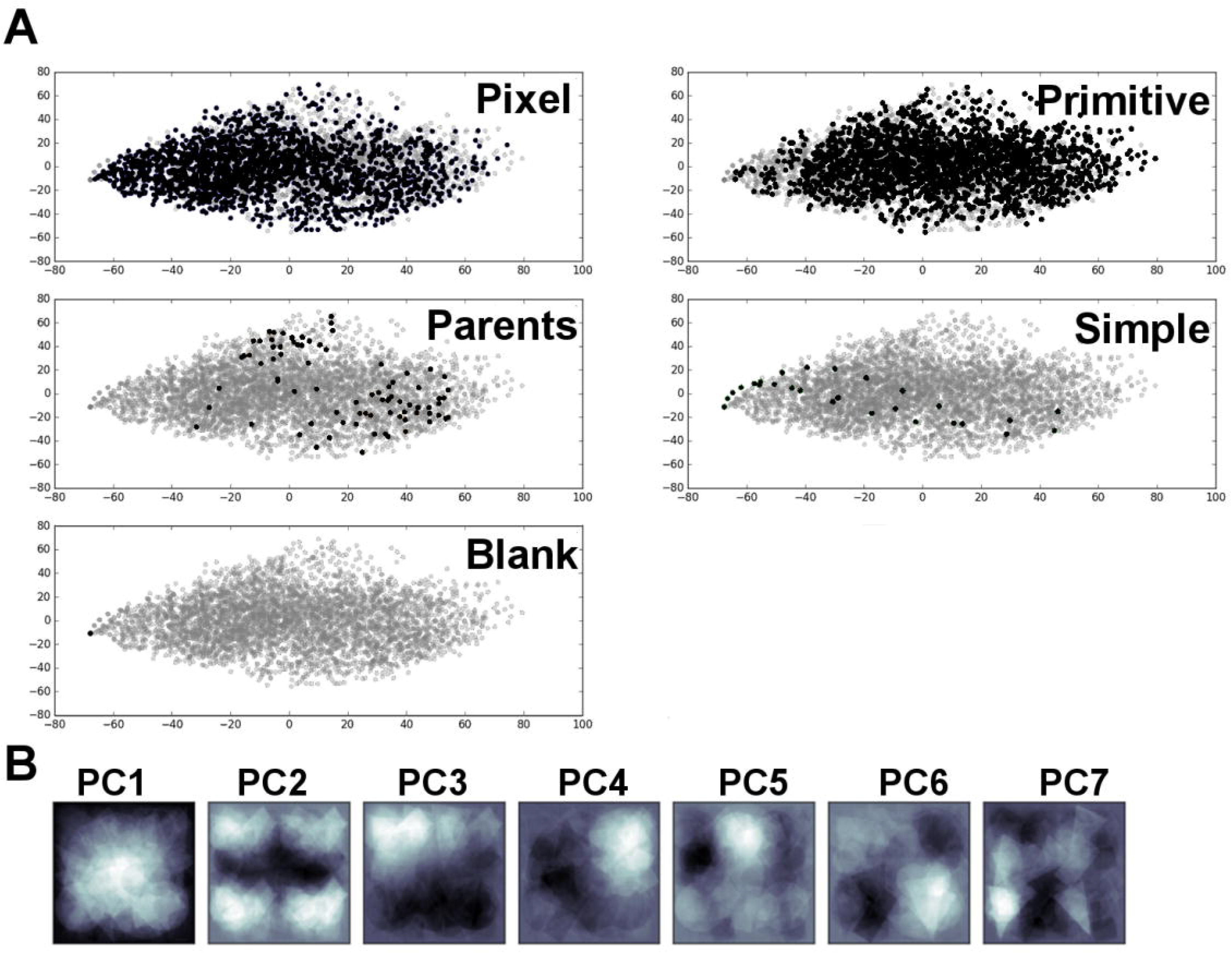
Principal Component Analysis of GTopoChip1. **A.** Scatterplot that represents every surface in first 2 Principal Components, split by groups. **B.** Image reconstruction of first 7 Principal components. White colour corresponds to highest variance.

We next visualized the first 7 PCs, which are called eigenimages (eigenvectors) (Figure 6B). The approach is broadly used in face recognition and is called eigenface generation ^25^. Design of every surface can be reconstructed by combining these eigenimages multiplied by their weights, which are unique for every topography. This visualization shows the most typical components of the surface design, specifically, where pixels (topography/non-topography) varied most between topographical features. The white color represents the highest variance and black — the lowest. Based on the first image (PC1), the component with the highest variance, it is clear that most variance between surfaces lies exactly in the centre of the topographical feature, confirming that coordinates of the primitives were sampled from the Normal distribution. PC2, second most important component maps to four repeated parts at the corners of the image, likely representing the original 10 μm features upscaled by the 2×2 format, and allowing permutation with 20 μm size features.

Other PCs represent patterns that occupy other parts of the image corners, and some patterns can also be traced back to the parent topographies. Interestingly, PC3 and PC4 represent similar patterns that are rotationally related. From a biological point of view, they represent exactly the same design because the cells will interpret these structures as identical. Such repeated designs were formed because our similarity comparison approach, based on the correlation index, did not take into account rotated versions of surface designs. While the rotation-agnostic similarity comparison test will decrease the redundancy of the designs, it will slow down the computational simulation. Another computationally expensive source of the redundancy is XY-offset of design elements. Implementation of fast surface design comparison test that accounts for rotation and XY-offset is a topic for further research.

### Comparison of GTopoChip1 and original TopoChip

The 81 parent topographies, used to seed the evolutionary process that generated GTopoChip1 were a subset of the 2176 unique topographies on the original TopoChip. We queried whether the GTopoChip1 is substantially different from the original TopoChip. To assess this, we summed all pixel values from the topographies of corresponding groups and generated a matrix. Figure 7A shows the matrix plotted using colours to represent the number of surfaces that have a pattern at the corresponding x y coordinate in the 2D image. For the original TopoChip, the surface was very smooth and consisted of 4 maxima because the original TopoChip consists of unique patterns of either 100×100, 200×200 or 280×280 pixels. We multiplied every 100×100 pattern 4 times to create a pattern with size 200×200. 280 × 280 patterns were excluded from our analysis as cropping them to the size of 200×200 would alter initial design significantly as not only shape of the pillars is important but also spacing between them.

**Figure 7.**
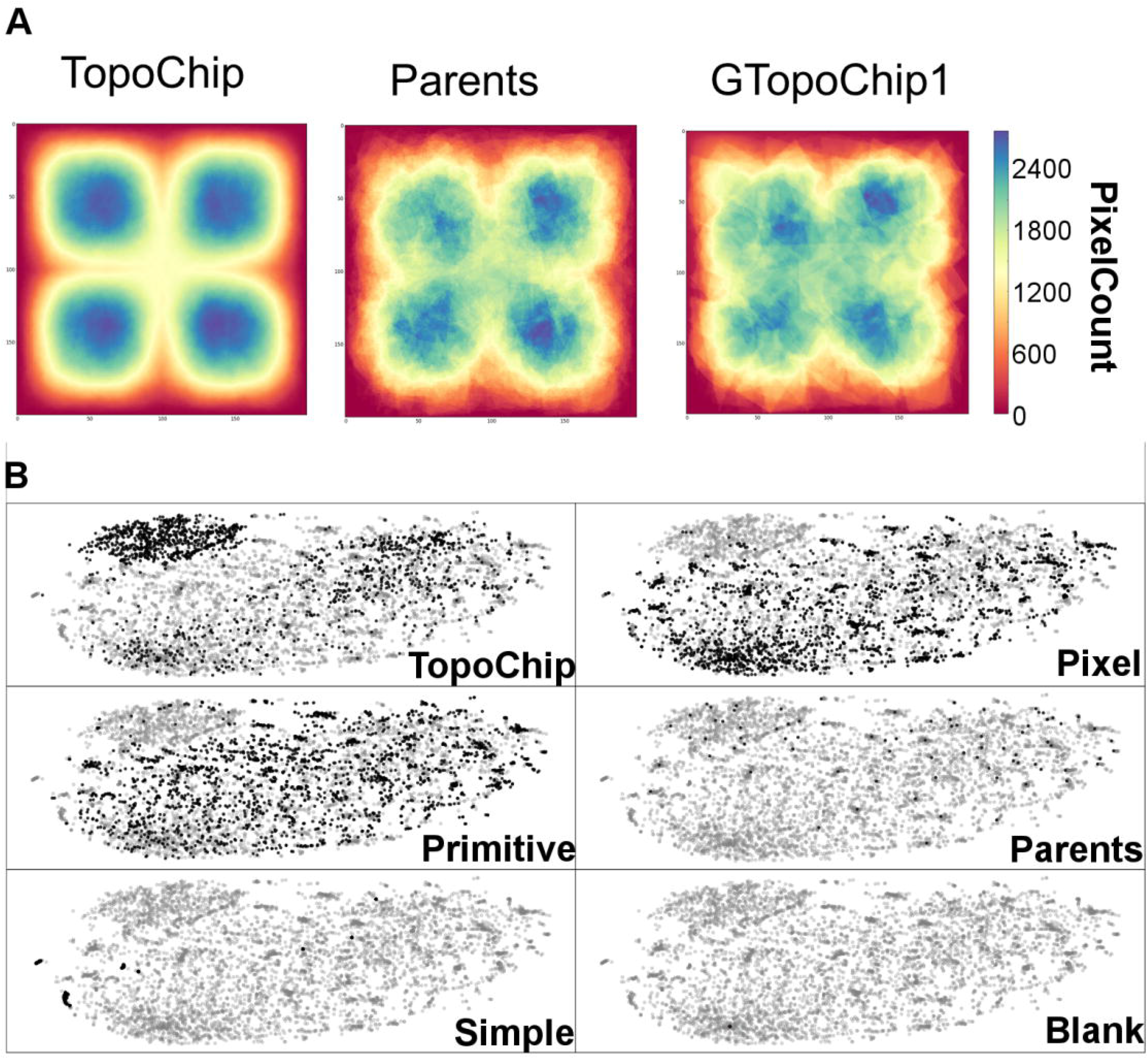
Comparison of GTopoChip1 and original TopoChip. **A.** Summation of all topographies pixels per group. **B.** t-SNE visualization of surfaces from TopoChip and GTopoChip1.

We used the t-SNE visualization algorithm, which applies nonlinear transformations to map multidimensional relations in a 2D plot, to compare these multidimensional data. t-SNE visualization was performed on the first 5000 PCs computed on images of the topographical features (pixel-based representation). Figure 7B clearly shows that some topographies from the original chip create a distinct cluster. Interestingly, small features occupy the left lower corner of the plot, and there is a large cluster at the top right corner of the plot (data not shown). The blank surface is in the lower left corner of the plot. These topographies generally do not induce high levels of ALP. Interestingly, the t-SNE plot created similar clusters regardless of whether pixel or feature based genomes were used (data not shown). Both topographies created by pixel and pattern design spanned homogeneously across the plot. However, some topographies created very dense areas, enriched with pixel-based topographies and sparse with primitive-based representations, and not represented by parents.

## Conclusion

The vastness of biomaterial design space precludes exploration by even the most optimistic brute force screening approaches. Although high throughput experimental methods clearly search for optimal materials more effeciently than conventional one-at-a-time experiments, they cannot explore more than a minute fraction of possibilities. However, evolutionary methods using iterative cycles of fitness assessment, mutation and selection are vastly more efficient in exploring these large parameter spaces as we have illustrated using the surface topography example. Genetic algorithms are driven by a fitness function describing the relationship between materials properties (representing a genotype) and function (representing the phenotype). AS an initial proof of concept, we studied ALP expression as a function of topography using experimental data. Our surface fabrication approach is efficient, as it takes the same effort to fabricate ten or thousands of surface topographies. We created a relatively large topographical library that allowed exploration of the topography/ALP expression relationship and generated the GTopoChip1 design.

Image-based representation of topographies generated drastically different topographies during evolution relative to the starting topographies. As the method cycled through, it generated new topographical features covering more design space compared to the parent surfaces and first-generation topographies. Clearly, imaged-based gene representations designs are not useful for evolving other biomaterial properties such as surface chemistry, or physicochemical properties such as softness, resilience etc. In these cases, more traditional chemical ‘genes’ can be used e.g. chemical graphs, text representations such as SMILES. As it will undoubtedly be useful to optimize topography and chemistry simultaneously, merging topographical and chemical genes into a “universal” gene representation is a topic that merits further investigated.

Future work will involve *in vitro* biological validation of the fabricated surfaces and the use of evolutionary methods to optimize topographies for more useful or realistic biological end points such as cell attachment, proliferation, and differentiation fate. Additionally, finding a surface inducing higher ALP expression, will validate the utility of the evolutionary approach, further elucidate the relationship between materials topographical properties and ALP expression, and allow further refinement of the algorithm.

## Material and Methods

### Algorithms

The algorithm for genetic perturbation of topographies was implemented in Python 2.7. We used the DEAP version 0.9a package for genetic algorithm calculations. Data and file manipulations were performed with packages Numpy, Json, pandas, scipy, and math. Figures were plotted with matplotlib and mayavi packages. Image manipulation was performed with the PIL, Codecs, cv2 and skimage packages. Surfaces identified as parents were selected by ALP expression greater than a threshold value. This threshold value was calculated as the 95 percentile of the ALP mean integrated intensities from all the surfaces. Topographical features (unique elements of the topography design) in the original chip had three sizes 10, 20 and 28 um. To permute designs with different sizes we excluded surfaces with size 28 microns and upscaled 10 um by merging four its copies in 2 × 2 format. Topographical feature design was obtained from the original design file that contains information about the type of the primitive, its size and location. For primitive-based parents, we used this data with no change, for the pixel-based approach we converted it to the 2D image with a size of 200 × 200 pixels, where pixels that correspond to pillars were set to one and no pillars to zero. For conversion into pixel-based genes, surface images were flattened to the one-dimensional matrix. The similarity between different surfaces was computed by Pearson correlation which was performed directly on flattened matrices of surface images. Total similarity per surface was computed by summing up absolute values of all its similarity indexes to other surfaces. Selection of algorithms for every step was done in random fashion. Generation of the topographies was done in multiple cycles as described in the main text. When all surfaces were generated they were shuffled in a random fashion. To meet with requirements of fabrication procedure all pillars with a diameter less than 4 μm were excluded from the final design.

## Acknowledgements

The research leading to these results has received funding from the European Union’s Seventh Framework Programme (FP7/2007-2013) under grant agreement no 289720. AV, AC and JdB acknowledge the financial contribution of the Province of Limburg. AC gratefully acknowledges her VENI grant (number 15057) from the Dutch Science Foundation (NWO).

## Author Contributions Statement

A.V., S.S., and A.C., performed the analysis. A.V., D.W., and J.B. wrote the main manuscript text.

A.V prepared figures 1-7. All authors reviewed the manuscript.

